# Denisovan admixture facilitated environmental adaptation in Papua New Guinean populations

**DOI:** 10.1101/2024.01.14.575483

**Authors:** Danat Yermakovich, Mathilde André, Nicolas Brucato, Jason Kariwiga, Matthew Leavesley, Vasili Pankratov, Mayukh Mondal, François-Xavier Ricaut, Michael Dannemann

## Abstract

Neandertals and Denisovans, having inhabited distinct regions in Eurasia and possibly Oceania for over 200,000 years, experienced ample time to adapt to diverse environmental challenges these regions presented. Among present-day human populations, Papua New Guineans (PNG) stand out as one of the few carrying substantial amounts of both Neandertal and Denisovan DNA, a result of past admixture events with these archaic human groups. This study investigates the distribution of introgressed Denisovan and Neandertal DNA within two distinct PNG populations, residing in the highlands of Mt Wilhelm and the lowlands of Daru Island. These locations exhibit unique environmental features, some of which may parallel the challenges that archaic humans once confronted and adapted to. Our results show that Denisovan-like haplotypes exhibit increased levels of population differentiation between PNG highlanders and lowlanders. The highly differentiated haplotypes, more common among highlanders, reside in genomic areas linked to brain development genes. Conversely, those more frequent in lowlanders overlap with genes enriched in immune response processes. Furthermore, Denisovan-like haplotypes displayed pronounced signatures of diversification within the major histocompatibility complex. Our findings suggest that Denisovan DNA has provided a valuable source of genetic variation to PNG genomes that facilitated adaptive responses to environmental challenges.

## Introduction

The initial human settlement of New Guinea is estimated to have occurred by at least 50 thousand years ago^1,2^. Today, the distribution of PNG populations across the region is uneven, often occurring in areas characterized by significant environmental disparities^3,4^. These environmental challenges, such as exposure to high altitudes or region-specific pathogens, have been demonstrated to correlate with phenotypic variations among PNG populations inhabiting distinct environments^5^. These challenges have also been established as factors contributing to the emergence of local genetic adaptation signatures^6,7^. Moreover, PNG, like their counterparts in near and remote Oceanian, bear a significant share of approximately 3-4% Denisovan DNA, ranking among the highest proportions globally^8,9^. This is in addition to the ∽2% of Neandertal ancestry that are found in PNG and all present-day non-Africans^10,11^. Our understanding of particularly the functional aspects of Denisovan DNA in its carriers and its potential contribution to adaptive processes in populations, such as the PNG, remains limited. This restricted knowledge is attributable to a variety of challenges. Firstly, the scarcity of fossil fragments has hindered the ability to comprehensively reconstruct Denisovan physiology and their historical geographic distribution^12^. These aspects make it challenging to postulate hypotheses regarding their functions in present-day individuals. Some insights have been derived from the Denisovan genome sequence8, such as the presence of only one copy of the amylase gene, hinting at potential differences in starch digestion^13^. Additionally, attempts to predict Denisovan phenotypes from genomic data have provided new insights into their skeletal physiology^14^. Due to the limited fossil fragments record originating from only a handful of locations, predicting the precise geographical range of Denisovans presents another significant challenge. However, both the distribution of these fossils and the distribution of populations carrying Denisovan DNA collectively suggest a relatively broad habitat^12,15^. It is plausible that Denisovans may have inhabited various regions across the Asian continent, spanning from Siberia to Southeast Asia. The existing uncertainty about their geographical distribution further complicates the formulation of hypotheses regarding potential environmental adaptations of Denisovans. The interbreeding between modern humans and Denisovans has been suggested to occur in multiple waves involving distinct Denisovan populations^7,16–18^. These findings may imply the existence of diverse Denisovan groups with potentially differing genetic compositions, adding another layer of complexity to the interpretation of their phenotypic characteristics. Another challenge arises when attempting to annotate the phenotypic effects of Denisovan DNA in contemporary individuals. In the case of introgressed Neandertal DNA, the utilization of phenotypic association data played a pivotal role in studying its influence on modern humans and drawing potential insights into Neandertal biology^19–23^. However, unlike Neandertal DNA, which is present in populations with accessible GWAS data, Denisovan DNA lacks readily available association information from phenotype and expression cohorts, rendering such annotation approaches less practicable. Nonetheless, several studies have provided valuable insights into how Denisovan DNA has influenced phenotypic variation and facilitated adaptation in modern humans^24–30^. Notably, among them are well-documented instances where Denisovan DNA has been associated with adaptations related to high-altitude environments, metabolic processes, and immune responses. The environmental challenges encountered by PNG as they dispersed across various regions could potentially mirror the challenges that Denisovans themselves had to contend with. Analyzing the fate of Denisovan DNA within PNG populations residing in diverse environmental conditions may offer a pathway to assess the functional capacity of Denisovan DNA and unveil some of its adaptive potential.

## Results

### Identification of archaic DNA in PNG highlanders and low-landers

In this study, our objective was to evaluate the degree to which DNA inherited from past interbreeding with archaic humans has contributed to shaping local adaptation of two geographically distinct PNG populations, each confronting unique environmental challenges. To address this task, we characterized the genomic landscape of Neandertal and Denisovan DNA in the genomes of 74 PNG individuals residing in the lowlands of Daru Island and 54 individuals inhabiting the highlands of Mount Wilhelm (Supplementary Table 1, Methods)^6^. We reconstructed introgressed archaic haplotypes utilizing a previously established method to detect archaic DNA within contemporary populations^22^. This approach identifies archaic SNPs (aSNPs) within the genomes of present-day individuals. These aSNPs function as markers of introgressed variants, with distinctive attributes such as allele sharing signatures with Neandertals and Denisovans. Moreover, these aSNPs reside on haplotypes of a length that exceeds those of segments that result from incomplete lineage sorting (ILS) between modern and archaic humans. ILS segments can produce comparable allele-sharing patterns to those seen in introgressed haplotypes but are on average considerably older and shorter (Methods). When employing this methodology in the analysis of PNG genomes within our study, we identified 168,413 aSNPs (Supplementary Table 2). These aSNPs were associated with 10,432 unique core haplotypes spanning across 50.0 to 65.5 megabases (Mb) of diploid archaic DNA within the autosome and chromosome X of the analyzed PNG individuals.

### Replication of introgression map results with an alternative method

We were able to replicate a large fraction of our identified archaic haplotypes using another alternative method (HMMIX) designed for reconstructing archaic DNA in contemporary populations^31^. This alternative approach employs a hidden Markov model to identify genomic regions characterized by a high density of SNPs that are absent in an unadmixed outgroup population. A significant proportion of 92.0% of the archaic haplotypes we identified were also captured by the alternative approach (with a posterior probability exceeding 0.8; 96.0% with posterior probability > 0.5; Methods). Generally, the alternative method detected a notably higher count of haplotypes (total haplotypes in PNG with posterior probability > 0.8, HMMIX: 483,376; our method: 172,121, Supplementary Figure 1a). However, many HMMIX-specific haplotypes with a posterior probability larger than 0.8 were characterized by an absence of archaic SNPs (81.2% HMMIX-specific haplotypes carried no archaic SNPs), possibly due to the underlying method of HMMIX which doesn’t involve archaic genome information. These haplotypes also exhibited shorter lengths compared to the ones we observed and omitted because their lengths aligned with incomplete lineage sorting (Supplementary Figure 1b-c).

### Most likely archaic source of introgressed haplotypes

Next, our goal was to assign the introgressed haplotypes we identified to their most probable archaic origins. To achieve this, we assessed their sequence similarity in comparison to the genomic sequences of three high-coverage Neandertals from the Altai Mountains^10^, Vindija Cave^11^, and Chagyrskaya Cave^32^, as well as the Denisovan individual^8^. We observed that, on average, ∽61% of the haplotypes per individual displayed a closer sequence resemblance to Neandertals than to the Denisovan individual (Figure 1a). This result stands in stark contrast to the anticipated genome-wide estimates of Neanderthal ancestry (∽2%) and Denisovan ancestry (∽4%) within these populations^8,11^. Yet, this figure aligns with prior studies that have attributed this disparity to the substantial differences between the sequenced Denisovan and the introgressing Denisovan population^27,28^. To address this challenge and enhance the ancestry assignment accuracy, our study introduced a secondary criterion based on the geographical distribution of archaic haplotypes (Supplementary Information). Assuming that the vast majority of archaic haplotypes with genuine Neandertal origin were introduced to the ancestors of all non-Africans, we scanned for the presence of Neandertal-like haplotypes from PNG individuals in contemporary Eurasian individuals that match the aSNP content and genomic location (Methods, Figure 1a). Our findings revealed that roughly half of these Neandertal-like haplotypes were exclusively found in PNG. This proportion is notably striking because, if these haplotypes are indeed of Neandertal origin, their presence solely in PNG suggests two potential scenarios: either their absence in other non-African populations resulted from their removal in them or they stem from a distinct Neandertal admixture pulse that differs from that shared with other non-Africans. Lacking prior evidence supporting either of these occurrences, we regard the likelihood of these explanations as minimal. Instead, we hypothesize that a significant portion of these PNG-specific Neanderthal-like haplotypes might be misclassified and are actually of Denisovan ancestry. Supporting this hypothesis, these PNG-specific haplotypes exhibited a closer sequence relationship to the Denisovan individual (Figure 1b) and displayed greater sequence divergence from Neandertals (Figure 1c) compared to the PNG Neandertal-like haplotypes identified in Eurasians. In our study we therefore decided to combine Denisovan-like haplotypes with the fraction of Neandertal-like haplotypes absent in Eurasian populations. The resulting new set of Denisovan-like haplotype accounted for 69.6% of all archaic haplotypes - a number substantially closer to the expected figure derived from allele-sharing estimates (Supplementary Information). Our analysis revealed that PNG highlanders and lowlanders carry similar amounts of Neandertal-like DNA (P=0.98, Mann-Whitney-U test). However, notably, PNG highlanders exhibited ∽3% more Denisovan-like DNA relative to the number of their lowland counterparts (P=0.003, Figure 1d, Supplementary Table 1).

**Figure 1:**
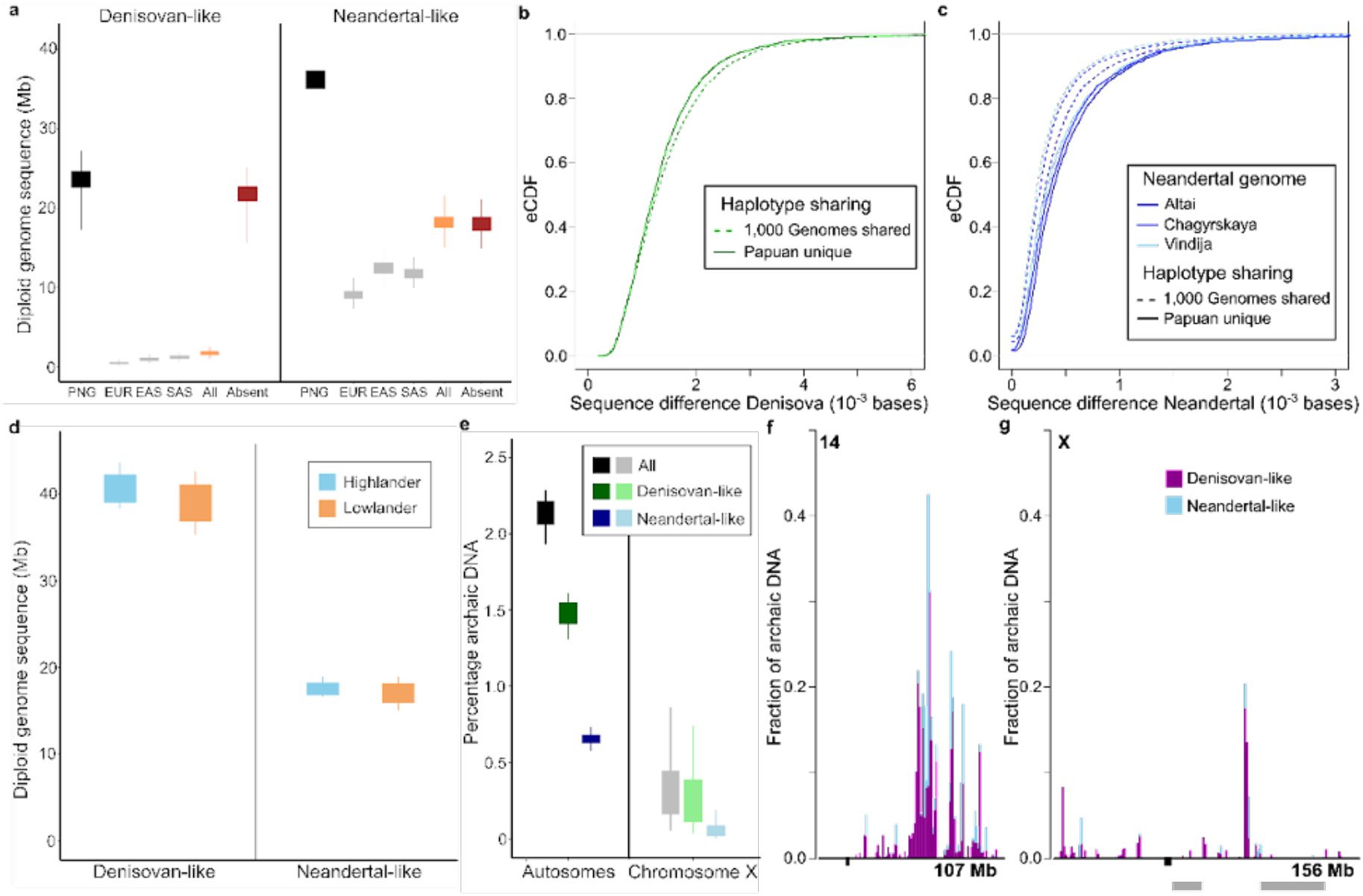
Archaic DNA in PNG genomes. **(a)** Amount of diploid genome sequences reconstructed in PNG individuals based on archaic haplotypes exhibiting higher sequence similarity with the Denisovan individual (left) and Neandertals (right). Boxplots are employed to visualize the distributions, with the outer whiskers indicating the minimum and maximum values. The recovered sequence amounts are presented in megabases (Mb) for various categories, including total sequences (black), sequences found in 1,000 Genomes Europeans (EUR, gray), East Asians (EAS, gray), South Asians (SAS, gray), and the combined sequences from all three Eurasian populations (All, orange). Additionally, the amount of genome sequences recovered using haplotypes that were not detected in any of the Eurasian populations (Absent) is displayed in red. **(b-c)** Empirical cumulative density distributions (eCDF, y-axis) are displayed for the sequence similarity measures (in number of differences per 10-3 bases, x-axis) of archaic haplotypes in PNG, specifically those exhibiting a greater sequence similarity with Neandertals than with the Denisovan individual. Panels (b) and (c) show these distributions in comparison to the Denisovan and three Neandertals, respectively. Haplotypes found in 1,000 Genomes Eurasians are represented by dashed lines, while PNG-specific haplotypes are delineated with solid lines. **(d)** Boxplots illustrating the amount of Denisovan-like and Neandertal-like diploid genome sequence in PNG highlanders (blue) and lowlanders (orange). Outer whiskers represent the minimum and maximum values. Unlike panel (a), where archaic ancestries are determined solely by sequence similarity, here, the annotation includes geographic refinement along-side sequence similarity. **(e)** Boxplots displaying the percentage of archaic (black/gray), Denisovan-like (green) and Neandertal-like (blue) ancestry on the autosomes and X chromosome in the PNG cohort. The outer whiskers indicate the 95% confidence interval borders. **(f-g)** The proportion of Denisovan-like (purple) and Neandertal-like (blue) DNA within one-megabase windows is depicted for chromosome 14 (f), which harbors the highest levels of archaic DNA, and chromosome X (g), characterized by the least amount of archaic DNA. Gray areas below the x-axis denote regions that were previously reported to be devoid of archaic ancestry in present-day populations.

### Genomic distribution of archaic haplotypes

Introgressed segments combined from all studied PNG individuals covered approximately 21.15% of the human genome. Among those segments we identified 21 archaic haplotypes characterized by archaic allele frequencies surpassing 70% in the combined dataset of the two PNG populations. Interestingly, only eight of these haplotypes exhibit overlaps with protein-coding genes, posing a challenge in establishing a functional link of those haplotypes on the biology of their carriers (Supplementary Table 3). Among the top five haplotypes featuring the highest archaic allele frequencies, only the Neandertal-like haplotype with the highest frequency of 89% (chr9:112,041,006-112,068,271) shares an overlap with a protein coding gene, namely *SUSD1*. Notably, *SUSD1* remains relatively understudied, although a recent investigation has suggested its involvement in neurodevelopmental disorders^33^. Additionally, we studied the archaic DNA content within regions previously identified as potential sites for negative selection acting on introgressed DNA^27,28^. Our analysis confirmed that, within all five autosomal genomic regions previously reported to be devoid of archaic DNA, there was either an absence or only negligible traces of archaic DNA present (Supplementary Figure 2). Consistent with these previous studies we also noted substantially lower levels of archaic DNA on the X chromosome (Supplementary Figure 2). PNG individuals exhibited an average 7.8-fold decrease in archaic DNA content on chromosome X in comparison to autosomes. This reduction was more prominent for Neandertal-like DNA, with a 12.6-fold decrease, compared to Denisovan-like DNA, which showed a 6.9-fold reduction (Figure 1e-g).

### Signatures of local adaptation in PNG lowlanders and highlanders

Next, we aimed to investigate the extent of population differentiation between the PNG highlanders and lowlanders. This analysis was designed to provide us with insights into the potential impact of selection on archaic haplotypes within each of these two examined populations. To achieve this, we conducted a comparative analysis of the allele frequency difference for each distinct archaic haplotype within both the PNG highlander and lowlander populations by calculating their respective Fst values (Methods, Figure 2a, Supplementary Table 3). We then determined whether archaic haplotypes exhibited altered levels of population differentiation between these two examined groups by comparing their Fst distribution to those of a genomic background of 1,000 randomly generated sets of non-archaic variants, chosen to match the allele frequency distribution of the archaic haplotypes (Methods, Figure 2b). Overall, our analysis revealed that the mean Fst value for archaic haplotypes (0.0178) was significantly lower than the mean values observed in the random non-archaic sets (ranging from 0.018 to 0.020, P<0.001). This trend was consistent for both Neandertal-like (P=0.02) and Denisovan-like (P<0.001) haplotypes. Next, we partitioned the Fst distribution according to whether the higher archaic allele frequency was detected in highlanders or lowlanders. The archaic Fst distributions consistently exhibited significantly lower mean values for haplotypes found at higher frequencies in lowlanders (P<0.001 for all haplotypes, including Denisovan-like and Neandertal-like haplotypes). However, we observed a noteworthy difference of the Fst distribution of archaic haplotypes with higher frequencies in highlanders. The Neandertal-like, Denisovan-like, and the combined Fst distributions exhibited higher mean values in comparison to the median of the Fst mean values from the random background sets. Notably, for the Denisovan-like haplotypes, this difference approached statistical significance (P=0.08) and was even more pronounced for the Denisovan-like haplotypes with an closer sequences similarity to the Denisovan than to any Neandertal (P=0.02). When we analyzed the Fst distribution of Denisovan-like haplotypes against randomly selected non-archaic background sets, we found that the top 10% of the Fst values for Denisovan-like haplotypes were significantly higher than those observed in the random non-archaic sets. (P<0.05, Figure 2c). This result suggests that particularly highly differentiated Denisovan-like haplotypes that show a higher frequency in highlanders show larger levels of population-differentiation than random frequency-matched non-archaic variants. Next, to further investigate signatures of selection on archaic haplotypes in PNG, we employed a computational approach to reconstruct the allele frequency trajectories of all archaic haplotypes over the course of the past 1,000 generations in both highlanders and lowlanders, utilizing an approximate full-likelihood method (Methods, Supplementary Table 4). We found that log-likelihood ratios (logLRs) that signal deviations from neutral allele frequency patterns showed a significant correlation with Fst values for both Denisovan-like (Spearman correlation, P=0.006) and Neandertal-like (P=0.03) haplotypes among highlanders, but this correlation was absent among lowlanders (P>0.05, Figure 2d). Our results indicate that the differences in frequencies among highlanders for some of the most distinct archaic haplotypes have been driven by substantial increases in allele frequencies over the past 1,000 generations, matching the time of between ∽25,000 and 20,000 years ago when highlands were settled permanently^4,34^. Our analysis of the reconstructed allele frequencies within highlander populations for the top 10 Denisovan-like and Neandertal-like Fst haplotypes revealed an average increase of 20% and 28%, respectively. Notably, there was a surge of 5.1% within the last 140 generations among the top ten Fst Denisovan-like haplotypes (Figure 2e). Comparatively, the average frequency increase for the top 10 Fst Denisovan-like and Neandertal-like haplotypes over the last 1,000 generations was notably smaller for lowlander populations, at 12% and 17%, respectively (Figure 2f). In general, our analysis revealed that many of the highly differentiated archaic haplotypes, particularly in PNG highlanders, were associated with specific allele frequency changes. The allele frequency burst of some of the most differentiated Denisovan-like haplotype within the last 100-200 generations suggest that recent selective pressures might have played a pivotal role in shaping the patterns of population differentiation we observed.

**Figure 2:**
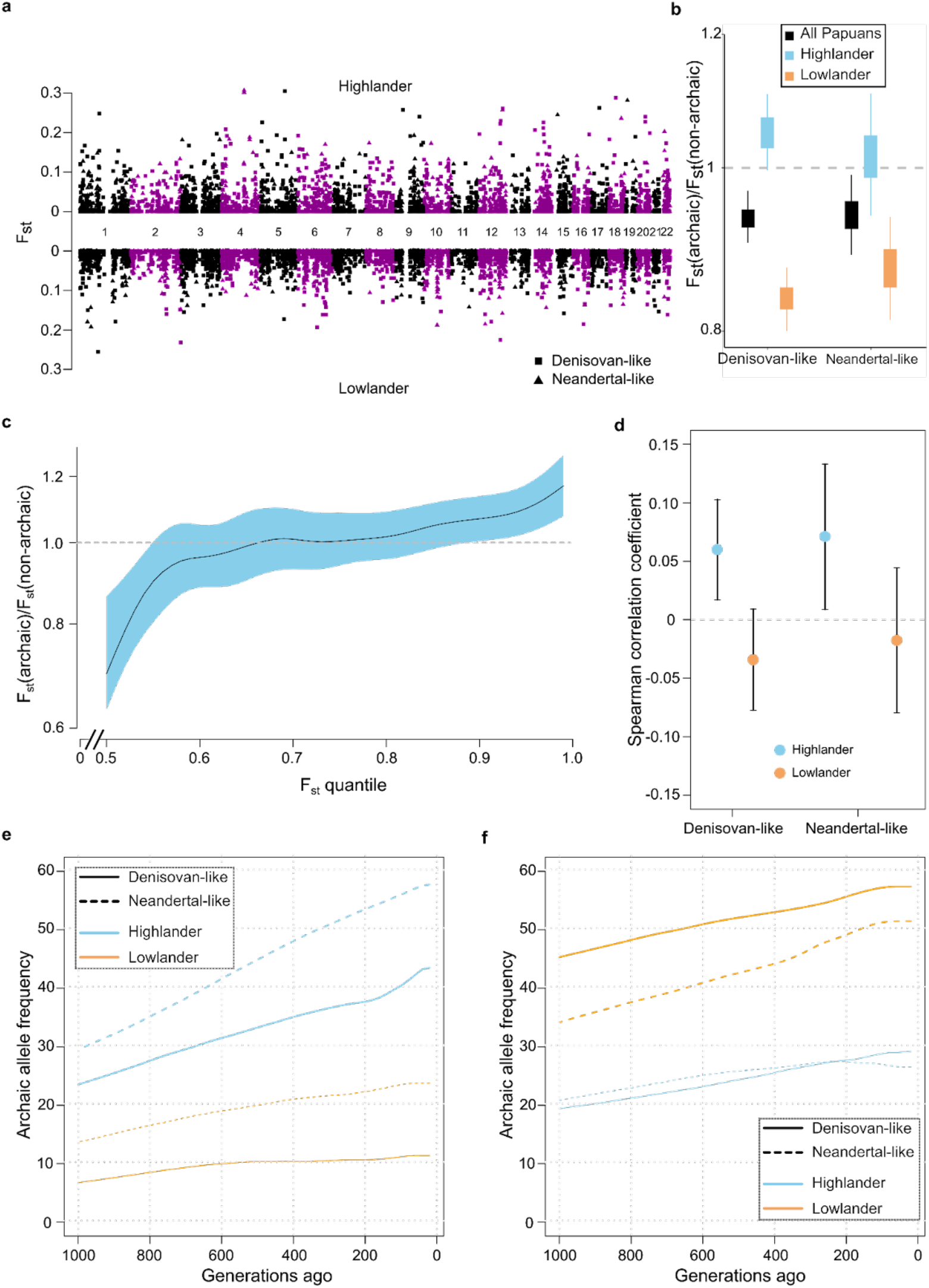
Frequency differences of archaic haplotypes between PNG highlanders and lowlanders. **(a)** Manhattan plot visualizes Fst values, representing frequency differences between PNG highlanders and lowlanders for Denisovan-like haplotypes (squares) and Neandertal-like haplotypes (triangles). The Manhattan plot is symmetrically divided across the x-axis, with data points displayed on both the upper and lower sides conditioned on whether a haplotype is found at a higher frequency in highlanders or lowlanders, respectively. **(b)** Distributions of the ratio of the mean Fst value for archaic haplotypes compared to each mean Fst value obtained from 1,000 non-archaic background sets (Methods). These distributions are presented for distinct categories of archaic haplotypes: the full sets of Denisovan-like (left, black) and Neandertal-like (right, black) haplotypes. Additionally, they include subsets specific to highlanders (blue) and lowlanders (orange) for both Denisovan-like and Neandertal-like haplotypes. Whiskers in the boxplots represent the 95% confidence interval for these distributions. **(c)** Distribution of the ratio between Fst quantiles for Denisovan-like haplotypes displaying a higher frequency in PNG highlanders and the quantile Fst values derived from 1,000 matching non-archaic background sets. The blue shaded area signifies the 95% confidence intervals, and the median ratio across quantiles is denoted by a black line. A gray line is included to represent the neutral expectation of one. **(d)** Spearman’s correlation coefficients together with their 95% confidence intervals (y-axis) calculated between Fst values and log-likelihood ratios calculated from reconstructed allele frequencies of Denisovan-like and Neandertal-like haplotypes are shown. Haplotypes are categorized into two sets: those exhibiting a larger archaic allele frequency in PNG highlanders (blue) and in lowlanders (orange). **(e-f)** Reconstructed average archaic allele frequencies (y-axis) over the past 1,000 generations for two sets of the top ten Denisovan-like (solid line) and Neandertal-like (dashed line) haplotypes with the highest Fst values in PNG highlanders (e) and lowlanders (f). The allele frequency trajectories are individually presented for each PNG population (highlanders: blue; lowlanders: orange).

### Gene content of genomic regions overlapping highly differentiated archaic haplotypes

A key emphasis of this study was to explore the potential phenotypic impact of archaic haplotypes exhibiting signs of local adaptation in highlanders or lowlanders. To tackle this question, we evaluated the gene content within genomic regions containing these haplotypes. Subsequently, to explore potential phenotypic consequences of genes within these regions, we conducted tests for functional enrichment within the gene ontology (GO)^35^. In total, we carried out four enrichment analyses for genes that overlapped with archaic haplotypes falling within the top 1% of the Fst distributions for each possible combination of higher archaic allele frequency in highlanders or lowlanders with Neandertal-like or Denisovan-like haplotypes (Supplemental Information, Methods, Supplementary Table 5). No enriched GO category was observed for high-Fst Denisovan-like haplotypes with higher frequencies in highlanders (FWER>0.05, Supplementary Table 5). Nevertheless, we observed a notable concentration of brain-related genes, including pivotal developmental genes like *NEUROD2* and *PAX5*, overlapping these haplotypes (Supplementary Information). This finding aligns with the discovery that among the four GO categories exhibiting the lowest P values, three were associated with fear response. This association might offer a potential target phenotype linked to these highlander-specific and highly differentiated Denisovan-like haplotypes. Genes located in genomic regions overlapping their counterparts, specifically high-Fst Denisovan-like haplotypes displaying a higher frequency in lowlanders, exhibited a GO enrichment for the category ‘cellular response to organic substance’ (FWER=0.03), along with related categories showing borderline significance linked to cytokine and protozoan responses (FWER: 0.06-0.08, Supplementary Table 5). Notably, among the Denisovan-like haplotypes associated with genes in these categories was a haplotype encompassing four members of the Guanylate-binding proteins family (*GBP1, GBP2, GBP4, GBP7*) - proteins crucial in immune response mechanisms^36^. This specific haplotype had been previously identified in Melanesians as a candidate for positive selection^6,27^ and carries an archaic variant introducing a missense variant to *GBP7* (Figure 3b, Supplementary Information). Additionally, this region exhibits high diversity among archaic humans, suggesting it might have been a target of admixture among these groups as well (Figure 3c). Among the two groups of high-Fst Neanderthal-like haplotypes, only the set exhibiting higher frequencies in highlanders displayed enriched GO categories. These categories comprised 22 associations related to regulatory and metabolic functions, largely influenced by members of the zinc finger family located within highlander-specific and high-Fst Neanderthal-like haplotypes (Supplementary Table 5, Supplementary Information).

**Figure 3:**
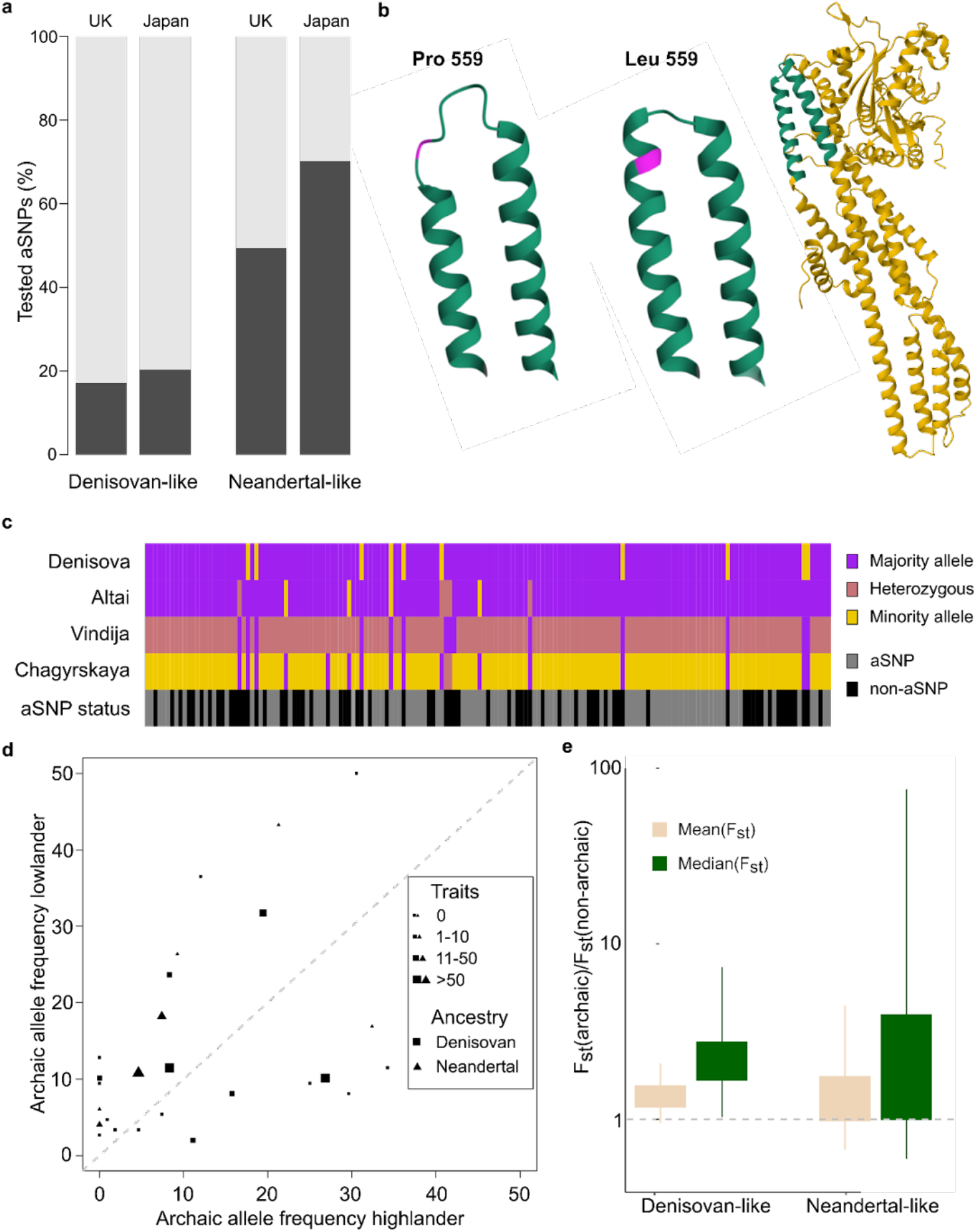
Phenotypic implications of archaic admixture in PNG populations. **(a)** The percentage of aSNPs tested in two biobank cohorts, namely the UK Biobank and Biobank Japan, is depicted. The relative distribution of aSNPs is categorized according to the ancestral origin of the haplotype linked to them in PNG. Only aSNPs with a minor allele frequency of at least 1% in both biobank cohorts are considered. **(b)** A three-dimensional representation of the human Guanylate Binding Protein 7 (GBP7) is depicted on the right. A distinct segment of the protein is emphasized in green. Two renditions of this protein segment are displayed: one reflects the modern human reference protein sequence (middle), while the other showcases an amino acid substitution introduced by the archaic allele rs139544379, resulting in a Leucine to Proline alteration at position 559 (left). **(c)** The genetic composition across 163 variable sites in the Denisovan and three Neandertal individuals within the region of the introgressed Denisovan-like haplotype at chr1:89,054,418.89,200,767 is depicted. Alleles are color-coded based on their prevalence in the four archaic genomes (major allele in purple, minor allele in yellow, and heterozygous sites in orange). The analysis only covers genetic positions with genotype information available for all four archaic individuals. The bottom panel highlights positions that represent aSNP locations in PNG (gray). **(d)** Scatterplot depicting the frequencies of archaic haplotypes spanning the Major Histocompatibility Complex (MHC) in PNG highlanders (x-axis) and lowlanders (y-axis). Denisovan-like haplotypes are represented by squares, while Neandertal haplotypes are denoted by triangles. The symbol sizes correspond to the number of trait associations associated with aSNPs segregating on these haplotypes (Methods). A gray diagonal line has been included to demarcate the region for haplotypes with identical frequencies in both populations. **(e)** Frequency distributions illustrate the ratio of both the mean (beige) and median (green) Fst value for archaic haplotypes overlapping the MHC compared to the mean and median Fst values obtained from 1,000 non-archaic background sets in the MHC (Methods). These distributions are presented for the sets of Denisovan-like (left, black) and Neandertal-like (right, black) haplotypes. Whiskers in the boxplots represent the 95% confidence interval for these distributions.

### Phenotypic inferences of archaic SNPs associated with highly differentiated archaic haplotypes

To better understand the potential impact of archaic DNA within the genomes of PNG, we conducted an analysis using data from UK Biobank^37^ and Biobank Japan^38^. This analysis aimed to identify associations involving aSNPs linked to archaic haplotypes reconstructed within the two PNG populations. It’s important to note that this approach has limitations due to notably lower levels of Denisovan DNA in UK and Japanese populations compared to PNG, resulting in a significantly reduced number of PNG aSNPs tested in these cohorts (Figure 3a). Moreover, this limitation was compounded by the possibility that aSNPs might have been tested on a different archaic haplotype in the biobanks compared to those in PNG. In total 379 archaic haplotypes harbored aSNPs exhibiting significant phenotype associations in at least one of the two biobank cohorts (Supplementary Table 6). Among these associations were 48 archaic haplotypes carrying aSNPs that alter the protein sequence of a gene. In general, we observed that a higher percentage of all Denisovan-like haplotypes carried missense aSNPs (5.9%) compared to Neanderthal-like haplotypes (4.7%, P=0.019, Fisher’s exact test, Supplementary Table 7). A total of 125 of all 568 missens-carrying haplotypes contained multiple missense aSNPs, notably a Denisovan-like haplotype carrying 10 missense aSNPs. This specific haplotype had been previously noted in a region involving *MUC19*, which has shown signs of introgression and selection among both archaic and modern humans^39^ (Supplementary Information). Traits commonly associated with these haplotypes included blood biomarkers, measurements related to bone density and body fat, as well as occurrences of diabetes. When we explored aSNP trait associations for all archaic PNG haplotypes, we identified four Denisovan-like haplotypes that exhibited substantial pleiotropy, displaying between 77 and 178 associations with various medical and non-disease phenotypes (Supplementary Table 6, Figure 3d). Intriguingly, all of these haplotypes were situated within the Major Histocompatibility Complex (MHC), a crucial immune-related region in the human genome. Notably, two of these haplotypes ranked among the top 10% in the Fst distribution. In total, we found 19 Denisovan-like and 7 Neandertal-like haplotypes located within the MHC. Both Denisovan-like and Neandertal-like MHC haplotypes showed significantly larger Fst values compared to their non-MHC counterparts (Denisovan-like: P=1.1×^−4^, Neandertal-like: P=6.5×^−4^). This result is consistent with the general higher diversity in this genomic region. However, Denisovan-like haplotypes also showed elevated median (0.067, P=0.03) and mean (0.059, P=0.10) Fst levels compared to other sets of frequency-matched non-archaic SNPs within the MHC (Figure 3e). We did not detect such a difference when comparing the Fst distributions of Neandertal-like haplotypes to random sets of frequency-matched non-archaic MHC SNPs.

## Discussion

In this study, we investigated the genomic and phenotypic impact of Denisovan and Neandertal DNA within two PNG populations living in distinct environmental regions - the mountainous terrain surrounding Mount Wilhelms and Daru Island. We found that Denisovan-like haplotypes exhibiting the most significant frequency differentiation between both populations exceeded the level of population differentiation seen in non-archaic variants with comparable frequencies. This effect was particularly pronounced for Denisovan-like haplotypes that had risen to higher frequencies among highlanders. These findings imply that Denisovan DNA played a substantial role in adaptive processes for these populations. Highly differentiated haplotypes that exhibited higher frequencies in highlanders overlapped several genes associated with early brain development. This result might reflect adaptive patterns to highlander-specific environmental factors. For instance, high-altitude-induced hypoxia has been linked to adaptive changes in neurons^40^. Similarly, dietary variations resulting from differences in food availability have been demonstrated to profoundly affect brain development^41^. Notably, none of the PNG individuals examined in our study carried a previously described Denisovan-like haplotype associated with high-altitude adaptation^25^. Therefore, at this juncture, it remains challenging to predict the extent and manner in which introgressed Denisovan DNA impacts these genes and their associated phenotypes. In this context it is worth noting that Neandertal DNA has been demonstrated to significantly impact several behavioral and neurological phenotypes in its carriers today^19,21,42–44^. Our findings imply that comparable patterns might also hold true for Denisovan DNA. Highly differentiated Denisovan-like haplotypes, present at high frequencies among lowlanders, exhibited a significant overlap with genes associated with pathogen response. One plausible explanation for this result could be the exposure to tropical diseases, such as malaria. A total of 94% of all malaria deaths in the Western Pacific region in 2021 were accounted for in Papua New Guinea^45,46^. Although malaria is highly prevalent in the geographical region, it is nearly absent among PNG highlanders^47^. Our findings imply that Denisovan DNA may have played a role in the adaptation to defense of malaria and/or other tropical diseases. Furthermore, we observed the presence of 19 Denisovan-like haplotypes within the Major Histocompatibility Complex, a critical immune region within the human genome. These haplotypes displayed significant frequency differences between highlanders and lowlanders, surpassing those found for frequency-matched non-archaic MHC variants. Our findings suggest that Denisovan admixture contributed to increased diversity in the MHC, a boost that might have been especially advantageous for the PNG population, which currently exhibits one of the lowest levels of heterozygosity worldwide^48^. Archaic admixture has frequently been acknowledged as a source of diversity, which is particularly relevant within the context of immunity and pathogen defense^49–54^. The influence of Denisovan-like haplotypes stands out when compared to introgressed Neandertal-like haplotypes. We do not observe a similar degree of population differentiation or functional significance for Neandertal DNA concerning population differentiation among PNG populations. However, we did discover that highly differentiated Neandertal-like haplotypes with greater prevalence among lowlanders were notably enriched for genes involved in transcriptional processes. These findings align with earlier reports highlighting the importance of Neandertal DNA in the regulation of gene expression^51,55–58^. Our findings offer potential phenotype-associated candidates that can contribute to a deeper understanding of the role of Denisovan DNA in contemporary adaptive processes. Furthermore, these candidates can serve as a basis for reconstructing the phenotypic characteristics of Denisovans and shedding light on the adaptive mechanisms within this archaic human group. These identified candidates present a valuable resource for functional testing through experimental assays^59,60^ and can collaborate with prediction algorithms to further explore their phenotypic significance^61^. Ultimately, a significant expansion of available association data^62^ will be a crucial component in advancing our understanding of the phenotypic potential of Denisovan DNA and its contribution to the adaptation of modern humans. It will also bring us another step closer to learning more about our extinct relatives and unique aspects of their biology.

## Acknowledgements

D.Y., M.A., V.P and M.D. were supported by the European Union through Horizon 2020 Research and Innovation Program under Grant No. 810645 and the European Union through the European Regional Development Fund Project No. MOBEC008. F.-X. R. and N.B. were supported by the French Ministry of Research grant Agence Nationale de la Recherche (ANR PAPUAEVOL 20-CE12-0003-01), the French Ministry of Foreign and European Affairs (French Prehistoric Mission in Papua New Guinea), and the Labex TULIP, France, and the Leakey foundation. Data analyses were carried out in part in the High-Performance Computing Center of the University of Tartu.

## Materials and Methods

### Genomic datasets

This study included high-coverage whole-genome sequencing data (version hg38) derived from a cohort of 128 unrelated adult PNG individuals, consisting of 54 individuals from the lowlands of Daru Island and 74 individuals from the highland region around Mount Wilhelm^6^. Furthermore, we analyzed high coverage whole-genome sequencing data from the 1,000 Genomes cohort^63^ and four archaic humans: the Altai, Chagyrskaya and Vindija Neandertals^10,11,32^ and the Denisovan^8^. Genotype information for all four archaic genomes was only available for the human genome version hg19. To harmonize our genotype datasets, genotypes were converted to hg38 coordinates using UCSC genome browser’s liftover tool^64^. In addition, genotypes were filtered using the provided genomic masks from each dataset. Genotype data from the 1,000 Genomes cohort was limited to single nucleotide variants (SNPs). Genotype data for the 1,000 Genomes cohort only contained variable positions within the dataset. All 1,000 Genomes individuals at genomic positions for which PNG or archaic humans showed at least one non-reference allele and that were without genotype information in the 1,000 Genomes cohort were considered to be homozygous for the human reference allele. Genotype information for the PNG and 1,000 Genomes individuals was available in phased format with the exception of chromosome X in PNG. We therefore processed the unphased genotype data for chromosome X using the same pipelines as previously used for the autosomes6. Briefly, we kept biallelic variants that had genotype information for more than 95% of individuals and carried the ‘PASS’ flag in the FILTER field. Next, we filtered out the pseudo-autosomal parts PAR1 and PAR2 and phased the remaining variants in the 128 PNG using shapeit4 with default parameters^65^.

### Archaic introgression map

We employed a previously established methodology^22^ to characterize introgressed archaic haplotypes in individuals of both PNG (N=128) and three 1,000 Genomes^63^ Eurasian superpopulations (Europe, N=633; South Asia, N=601, East Asia, N=585). This approach identifies these haplotypes by leveraging distinctive characteristics, including shared allele signatures, haplotype structure, and haplotype length, which are indicative of ancestral interbreeding between modern humans and Neandertals as well as Denisovans. Following the approach, we first identified archaic SNPs (aSNPs) within the genomes of four distinct populations under analysis: PNG, East Asians, South Asians, and Europeans. These aSNPs were defined as those containing an allele that met the following criteria: (i) it was absent in the 1,000 Genomes Yoruba population, (ii) it was present in at least one of the three high-coverage Neandertal genomes (Vindija^11^, Chagyrskaya^32^, Altai^10^), or the Denisovan genome8, and (iii) it was present in at least one individual within the four populations we were examining. Next, within each of the four populations, we computed pairwise measures of linkage disequilibrium (LD) represented as r^2^ between all identified aSNPs within that particular population. We collapsed sets of aSNPs that showed r^2^>0.8 and defined them as an archaic haplotype. All aSNPs displaying no LD of r^2^>0.8 with any other aSNP were removed. Subsequently, we calculated the nucleotide distance for all remaining haplotypes by measuring the span between the two furthest aSNPs within each haplotype. We then assessed the compatibility of each haplotype’s length with the genomic phenomenon incomplete lineage sorting (ILS). ILS refers to the retention of ancestral genetic variation shared by some modern and archaic human populations, which predates the divergence of these human groups. As a result, ILS can lead to similar allele-sharing patterns as aSNPs. However, because ILS segments are considerably older, the average length of ILS haplotypes is much shorter compared to Neandertal or Denisovan haplotypes. These archaic haplotypes typically span tens of kilobases in size, reflecting the relatively recent admixture between modern and archaic humans approximately 55,000 years ago^66^. Building upon the methodology introduced by Huerta-Sanchez et al^25^ and incorporating more recent estimates for divergence and mutation rates^50^, we computed the likelihood of each inferred haplotype’s length being compatible with incomplete lineage sorting (ILS). This calculation was conducted using recombination rate estimates obtained from two separate cohorts^67,68^. We calculated false-discovery rates (FDR) by adjusting the acquired P values for multiple testing using the Benjamini-Hochberg procedure^69^. Haplotypes that exhibited a length compatible with ILS for both recombination maps (FDR>0.05) or had fewer than 10 aSNPs were excluded from further analysis. Lastly, using the remaining haplotypes and their corresponding aSNPs, we reconstructed haplotypes within individuals across all investigated populations, based on each individual’s alignment with the archaic allele at the aSNP sites. Next, we assigned archaic haplotypes to their most probable archaic source by evaluating their sequence similarity with the genomes of three high-coverage Neandertals and the Denisovan. For haplotypes alleles that were present in a heterozygous state in the unphased archaic human genomes we defined a distance of 0.5. Haplotypes were defined as Denisovan-like when they exhibited a stronger sequence affinity with the Denisovan genome than with the other three Neandertals. All remaining haplotypes were categorized as Neandertal-like. Moreover, we further categorized Neandertal-like and Denisovan-like haplotypes within the PNG population based on their presence in any of the three Eurasian populations we examined. Archaic haplotypes in PNG were regarded as shared with Eurasian populations if they exhibited an overlap of at least 80% and were composed of the same set of aSNPs in the overlapping region as the matching archaic haplotypes in Eurasians.

### Evaluation of introgression map

To assess the performance of our approach, we leveraged HMMIX, an independent method utilizing a Hidden Markov model designed to infer putative introgressed segments^31^. We executed the method using default parameters for all 128 PNG samples. To closely mirror the comparison with our approach, we selected the Yoruba population from the 1,000 Genomes cohort as the outgroup. In addition, we used the four archaic genomes to annotate HMMIX’s results with shared archaic variants. In comparing our results to the output of HMMIX, we examined the overlap between fragments identified as “Archaic” by HMMIX and our inferred archaic haplotypes, considering overlaps of any length.

### Measures of population differentiation

We assessed the extent of population differentiation between PNG highlanders and lowlanders by calculating Fst values for all identified archaic haplotypes in these two groups. For each haplotype, we randomly selected candidate aSNPs for the analysis. Per haplotype, we conditioned the selected aSNP to be within the same 1% allele frequency bin as the aSNP with the median allele frequency value of a given archaic haplotype. We used VCFtools (0.1.14) software and computed the Weir and Cockerham Fst estimate^70^. Furthermore, we created 1,000 control sets of randomly selected non-archaic SNPs that matched the number of SNPs and their frequency distribution of the tested aSNPs within the combined PNG population dataset.

### Computational reconstruction of allele frequency trajectories for aSNPs

We employed computational methods to reconstruct allele frequency trajectories, utilizing genomic data from both PNG highlanders and lowlanders, applying a modification of the pipeline detailed in André et al^6^. In brief, our approach involved selecting three random representative aSNPs from each archaic haplotype. Subsequently, we extracted the local genealogical tree for each aSNP, utilizing the RELATE software^71^ (v1.1.8). These generated trees served as input for CLUES^72^, an approximate full-likelihood method for testing signatures of selection (v1). CLUES facilitated the reconstruction of allele frequencies and assigned log-likelihood ratios (log(LR)) to indicate support for non-neutrality. We did not assess aSNPs with a minor allele frequency below 5%. Furthermore, we excluded data points for which the CLUES algorithm did not yield a log(LR) value.

### Functional enrichment analysis

We performed functional enrichment analysis in the Gene Ontology (GO)^35^ using the R package GOfuncR^73^. Specifically, we examined four sets of archaic candidate haplotypes that ranked within the top 1 percentile of their respective Fst distributions. These four sets were created by pairing (i) highlanders or lowlanders with (ii) Denisovan-like haplotypes or Neandertal-like haplotypes. For each of the four sets, we defined background haplotypes, which included all remaining haplotypes that did not fall within the top 1 percentile of their respective Fst distributions. To account for the genomic clustering of functionally related gene groups, we ran the GO enrichment software with the parameter circ_chrom=TRUE. This setting allowed us to create test sets that randomly shifted the coordinates of both candidate and background haplotypes on the circularized version of the same chromosome. Family-wise error rates (FWER) were determined by comparing empirical enrichment P values for each GO category to the minimum enrichment P values observed in the entire GO for each test set.

### Phenotypic and regulatory annotation of aSNPs

We conducted a screening of phenotypic association data from two biobank cohorts, the UK Biobank^37^ (4,280 phenotypes, https://www.nealelab.is/uk-biobank) and Biobank Japan^38^ (220 phenotypes), to identify associations with aSNPs linked to the archaic haplotypes observed in PNG highlanders and lowlanders. We included in our analysis all associations involving aSNPs with allele frequencies exceeding 1% and meeting the stringent genome-wide significance threshold of P<5×^−8^. In addition, we leveraged ENSEMBL’s variant effect predictor74 (ensembl-vep version: 109.3) to annotate the aSNPs identified in PNG with their predicted molecular consequences.

### Protein visualization

We employed AlphaFold2^75^ to predict the 3D protein folding structures for two variants of GBP7. The first variant was constructed based on the reference protein sequence, while the second variant incorporated a Proline substitution in place of Leucine at amino acid position 559. This substitution was introduced by a Guanine on an aSNP at position chr1:89,132,390, with the reference allele being Adenine. This aSNP resides on the Denisovan-like haplotype (chr1:89,054,418-89,200,767), which encompasses this gene. To visualize the resulting protein structures for both variants, we utilized the Mol* viewer^76^ (Figure 3b).

