## Supplementary Information for "Denisovan admixture facilitated environmental adaptation in Papua New Guinean populations"

*Most likely archaic source of introgressed haplotypes*

We assigned introgressed haplotypes to their most probable archaic origins by quantifying their sequence similarity in comparison to the genomic sequences of three high-coverage Neandertals from the Altai Mountains^1^, Vindija Cave^2^, and Chagyrskaya Cave^3^, as well as the Denisovan^4^. In our initial categorization, haplotypes exhibiting the highest sequence similarity to the Denisovan genome were classified as Denisovan-like haplotypes, while all other haplotypes were categorized as Neandertal-like. Although the quantity of Neandertal-like and Denisovan-like haplotypes was similar in both of the studied PNG populations, the relative proportions of these two types of haplotypes did not align with previous estimates of Neandertal and Denisovan DNA in these populations (Figure 1a). Based on genome-wide estimates derived from allele-sharing statistics^5^, we would expect approximately ~2% Neandertal DNA and ~4% Denisovan DNA in the genomes of contemporary PNG^2,4^. Consequently, we anticipated that Denisovan-like haplotypes constitute around 2/3 of all introgressed haplotypes. Our observations reveal a contrasting scenario, with Neandertal-like haplotypes accounting for 60.7% of all identified haplotypes. Previous studies that have examined archaic DNA within PNG genomes and other populations featuring substantial Denisovan ancestry^6,7^ have described a similar pattern. They have attributed the diminished proportion of Denisovan-like DNA in Oceanian populations to the large divergence between the genome sequences of the Denisovan individual and the introgressing Denisovan population. In this study, we aimed to enhance the annotation of archaic haplotypes by integrating information regarding their global geographical distributions. Denisovan DNA has been documented in several Oceanian populations, with sporadic and limited occurrences in certain Asian populations^6–8^. Our hypothesis was that Denisovan-derived archaic haplotypes in PNG would be mostly absent in mainland Eurasian populations, providing a means to identify Denisovan haplotypes that cannot be distinguished solely based on sequence similarity. To investigate this, we examined the presence of archaic haplotypes in PNG within archaic haplotypes we reconstructed in the 1,000 Genomes Eurasian dataset^9,10^ (Methods). We found that only a small proportion of Denisovan-like haplotypes were identified in Eurasians (7.3%). In contrast, around half of the Neandertal-like haplotypes were not present in Eurasian populations (50.0%). Combining Denisovan-like haplotypes with the fraction of Neandertal-like haplotypes absent in Eurasian populations resulted in a proportion of 69.6% of all archaic haplotypes - a figure much closer to the expected number derived from allele-sharing estimates. Another indicator that underscores the effectiveness of this approach is the observation that PNG-specific Neandertal-like haplotypes display a greater sequence distance from Neandertals and a shorter distance to Denisovans when contrasted with Neandertal haplotypes in PNG that are shared with Eurasians (Figure 1b-c). These shared haplotypes exhibit a shorter distance to Neandertals (Mann-Whitney-U test, P=5.7x10^-101^) and a greater distance to Denisovans (Mann-Whitney-U test, P=0.005). Given the improved Neandertal-to-Denisovan ratio achieved through geographical refinement and the support provided by the sequence similarity analysis, we decided, for the remainder of this manuscript, to classify Neandertal-like haplotypes absent in Eurasians as Denisovan-like haplotypes as well. We acknowledge that by using this approach we will mistakenly categorize actual PNG-specific Neandertal haplotypes as Denisovan-like and fail to re-classify actual Denisovan haplotypes that show a higher sequence affinity to Neandertals if they are also present in Eurasians. Nonetheless, we anticipate this impact of both cases to be minimal, a hypothesis corroborated by the Neandertal-to-Denisovan ratio, which closely aligns with the anticipated figure.

*Gene content of genomic regions overlapping highly differentiated archaic haplotypes*

We first assessed the gene content within these genomic regions within haplotypes exhibiting signs of local adaptation in highlanders or lowlanders. We tested for functional enrichment within the gene ontology ^11^ for genes that intersected with archaic haplotypes with substantial population differentiation levels between highlanders and lowlanders. To accomplish this, we carried out four enrichment analyses for genes that overlapped with archaic haplotypes falling within the top 1% of the Fst distributions for each possible combination of higher archaic allele frequency in highlanders or lowlanders with Neandertal-like or Denisovan-like haplotypes (Methods, Table S5). Genes overlapping Denisovan-like haplotypes with higher frequencies in highlanders did not exhibit any significant GO enrichment (FWER>0.05). Notably, among the five categories displaying the most robust enrichment results (P<0.001), four of them were linked to the behavioral fear response. These four categories were significantly impacted by the presence of *NEUROD2* and *FBXL20*, both of which overlap with a haplotype located on chromosome 17. This particular haplotype is highly prevalent among highlanders, with a frequency of 49% (ranking as the third highest Fst value for Denisovan-like haplotypes), while its frequency among lowlanders is substantially lower at 13%. Both *FBXL20* and *NEUROD2* are expressed in neurons, with *NEUROD2* playing a pivotal role in neuronal development ^12,13^. Notably, among the 16 genes associated with two of the top five Fst Denisovan-like haplotypes in highlanders, 10 are closely linked to brain function. These genes encompass a range of functions related to brain and neuron development, including *PAX5**^14^*, *PGAP3**^15^* and *CDK12**^16^*, as well as *PNMT*, a key component in adrenaline production^17^, *MED1* associated with the circadian clock^18^, and *PPP1R1B*, a critical mediator in dopamine signaling pathways, influencing mood and motivation, among other functions^19^ (other brain-related genes: *TCAP**^20^*; *IKZF3**^21^*). Although a formal Gene Ontology (GO) enrichment analysis did not yield significant results, the notable concentration of brain-related genes within genomic regions that overlap Denisovan-like haplotypes exhibiting substantial frequency increases in PNG highlanders suggests a potential role for Denisovan DNA in influencing the biology of the brain in this population. However, we did observe an enrichment of genes associated with two of the remaining three enrichment analyses for groups of ancestry-specific high Fst haplotypes. More specifically, genes overlapping the top 1% of Denisovan-like Fst haplotypes, where the higher archaic allele frequency was found in lowlanders, were enriched for the Gene Ontology category 'cellular response to organic substance' (FWER=0.03). This category comprised 17 genes that overlapped with 12 of the tested high Fst haplotypes. Based on the reconstructed archaic allele frequencies for these haplotypes, we found that their average frequency steadily increased over the last 1,000 generation in lowlanders (from 42.2% to 56.9%), a pattern that was less pronounced in highlanders (from 23.2% to 28.8%). Furthermore, we observed three closely related and more specific Gene Ontology categories linked to cytokine and protozoan responses, which displayed comparable but borderline significant enrichment signatures (FWER: 0.06-0.08, Table S5) and were associated with subsets of these genes. Of particular significance were four members of the Guanylate-binding proteins family (*GBP1*, *GBP2*, *GBP4*, *GBP7*). This gene family encodes proteins that play a crucial role in the innate immune response. These proteins are involved in host defense against pathogens, particularly in the context of intracellular infections. GBPs can target and disrupt the membranes of intracellular bacteria and parasites, helping to control infections and modulate the immune response^22^. Three of these family members were associated with the Denisovan-like haplotype which exhibited the highest observed Fst value among all tested archaic haplotypes (chr1:89,054,418-89,200,767). The archaic haplotype is found at 89% in lowlanders and 54% in highlanders. The associated genomic region has previously already been identified as a candidate for positive selection in PNG lowlanders^23^ and has been associated with a high-frequency archaic haplotype in another Melanesian cohort^6^. One aSNP linked to the GBP haplotype we identified induces a change in the protein sequence of *GBP7*, specifically a Leu559Pro modification (Figure 3b), which has been predicted as deleterious and possibly damaging by two different prediction methods^24,25^. Notably, we discovered that the genomic region overlapping with this particular haplotype exhibited variability in the genomes of archaic humans. While the Altai Neandertal and the Denisovan individual showed high sequence similarity in that regions and shared most of the archaic alleles associated with the introgressed haplotype identified in PNG, the Vindija Neandertal was predominantly heterozygous in that region, and the Chagyrskaya Neandertal showed the largest sequence difference (Figure 3c). Another group of genes that displayed enrichment in the Gene Ontology were those that overlapped with high-Fst Neandertal-like haplotypes, where the higher archaic allele frequency was observed in highlanders. In total, we identified 22 categories with a family-wise error rate below 0.05. These categories exhibited close functional relationships and were primarily associated with transcriptional, regulatory and metabolic activities (Table S5). These enrichment patterns were strongly influenced by six members of the zinc finger family, one of the largest families of transcription factors and gene expression regulators in the human genome^26^.

*Phenotypic inferences of archaic SNPs associated with highly differentiated archaic haplotypes*

To enhance our phenotypic inferences, we conducted an examination of phenotype databases to identify associations involving the aSNPs we reconstructed within the two PNG populations. However, the databases we examined, namely the UK Biobank^27^ and Biobank Japan^28^, while providing association data for a wide range of phenotypes, are not ideal for this specific task. Firstly, it was observed that only a subset of PNG aSNPs were subjected to testing (Figure 3a, Table S6). Importantly, data on the associations of aSNPs carried on Denisovan-like haplotypes were notably absent. Additionally, due to the shared ancestry between Neandertals and Denisovans, it is likely that aSNPs located on Denisovan-like haplotypes with available association information were assessed within the context of a Neandertal-like haplotype background. Given the differing composition of aSNPs between Neandertal-like and Denisovan-like haplotypes, it remains uncertain whether the association findings from these two Eurasian cohorts can be extrapolated to PNG populations. Nonetheless, a previous study comparing a blood eQTL dataset from Indonesia with blood eQTLs from Eurasian cohorts^29^ showed that replication between these cohorts is feasible to a certain extent. Overall, we identified genome-wide significant associations in the two examined biobanks for aSNPs linked to 379 archaic haplotypes. As expected, Denisovan-like haplotypes less frequently carried aSNPs associated with phenotypes (Figure 3a). Several phenotype associations were identified in connection with aSNPs that altered the protein sequences of specific genes. In total, we observed that 757 aSNPs present on archaic haplotypes in PNG had a protein-sequence-altering impact on a total of 592 genes (Table S7). A higher proportion of Denisovan-like haplotypes (5.9%) than Neandertal-like haplotypes (4.7%) carried missense variants (P=0.019, Fisher’s exact test). Twenty-two percent of haplotypes carried more than one missense aSNP variant. A Denisovan-like haplotype on chromosome 12 (chr12:40,261,071-40,434,217, Allele frequency: highlander 7.4%, lowlander 10.1%) carried a total of 10 aSNPs that changed the protein sequence of Mucin 19 (*MUC19*). *MUC19* has previously been reported to overlap a genomic region which has been a target of introgression and selection among archaic and modern humans^30^. Notably, among the missense-carrying archaic haplotypes, 48 missense aSNP exhibited significant associations with phenotypes (Table S7). The predominant traits associated with these haplotypes included various blood biomarkers, along with measurements related to body fat and bone density. Additionally, we found that two Neandertal-like haplotypes harbored missense aSNPs associated with diabetes.

**Supplementary tables**

**Table S1: PNG sample information.** The table comprises details regarding the sex, sampling location, access to the European Genome-Phenome data repository, and the quantities of reconstructed Denisovan-like and Neandertal-like DNA found in the analyzed individuals from PNG in this study.

**Table S2: List of archaic SNPs (aSNP) reconstructed in PNG individuals.** The table presents a comprehensive list of aSNPs linked to Denisovan-like and Neanderthal-like haplotypes found in individuals from Papua New Guinea (PNG) within this study. It includes aSNPs located on autosomes and chromosome X, along with details about the allelic state in the Yoruba population and the high-coverage genomes of three Neanderthals and the Denisovan. In cases where the Yoruba state is unavailable (“./.”), we assumed the Yoruba population from the 1,000 Genomes Project to have a consistent allele state with the human reference allele. Moreover, the table incorporates data on the frequency of aSNPs within PNG individuals, along with information about the location and probable ancestry of the archaic haplotype they are associated with.

**Table S3: List of archaic haplotypes in PNG individuals.** The table presents a comprehensive list of all identified archaic haplotypes located on the autosomes and the X chromosomes of Papua New Guinea (PNG) individuals investigated in this study. It includes details regarding their frequency within the PNG populations, Fst estimates between highlander and lowlander groups in PNG, aSNP content, and the most likely archaic source population.

**Table S4: Selection inferences of archaic haplotypes in PNG populations.** The table shows summary statistics generated by the CLUES algorithm, indicating potential instances of recent selection on archaic haplotypes within both PNG highlander and lowlander populations. Each archaic haplotype underwent testing involving three associated aSNPs. For each aSNP the inferred selection coefficient alongside its corresponding log(LR) value are displayed. Instances where the archaic haplotype was not present in one population or the CLUES algorithm didn’t converge show ‘NA’ values.

**Table S5: Gene ontology enrichment for high Fst haplotypes.** The table exhibits summary statistics obtained from Gene Ontology (GO) enrichment analyses conducted on four gene sets. These sets comprised genes that overlapped with the highest 1% Fst Denisovan-like and Neandertal-like haplotypes within both highlander and lowlander populations in Papua New Guinea (PNG). Categories presented in this table include all those with a family-wise error rate (FWER) below 0.05. Additionally, it includes the top 10 categories characterized by the lowest P values despite having an FWER above 0.05. Moreover, the table showcases all genes associated with both the tested haplotype set and the particular category under investigation.

**Table S6: Phenotype associations of archaic haplotypes in PNG populations.** The table includes a list of archaic haplotypes containing aSNPs that exhibited genome-wide significant associations in either the UK Biobank or Biobank Japan. Additionally, it presents a compilation of associated traits. We annotated haplotypes that either bore missense variants or were situated within the Major Histocompatibility Complex.

**Table S7: Predicted molecular effects of aSNPs in PNG populations.** The table comprises a list of all aSNPs detected in PNG populations included in this study with predicted molecular effects, according to ENSEMBL’s variant effect predictor, that either have a ‘HIGH’ predicted impact or fall into the categories ‘missense_variant’, ‘3_prime_UTR_variant’, ‘5_prime_UTR_variant’ or TF_binding_site_variant’. The table also presents the estimated level of deleteriousness for aSNPs predicted to modify a gene's protein sequence, as determined by the PolyPhen2 and SIFT algorithms.

**Supplementary figures**

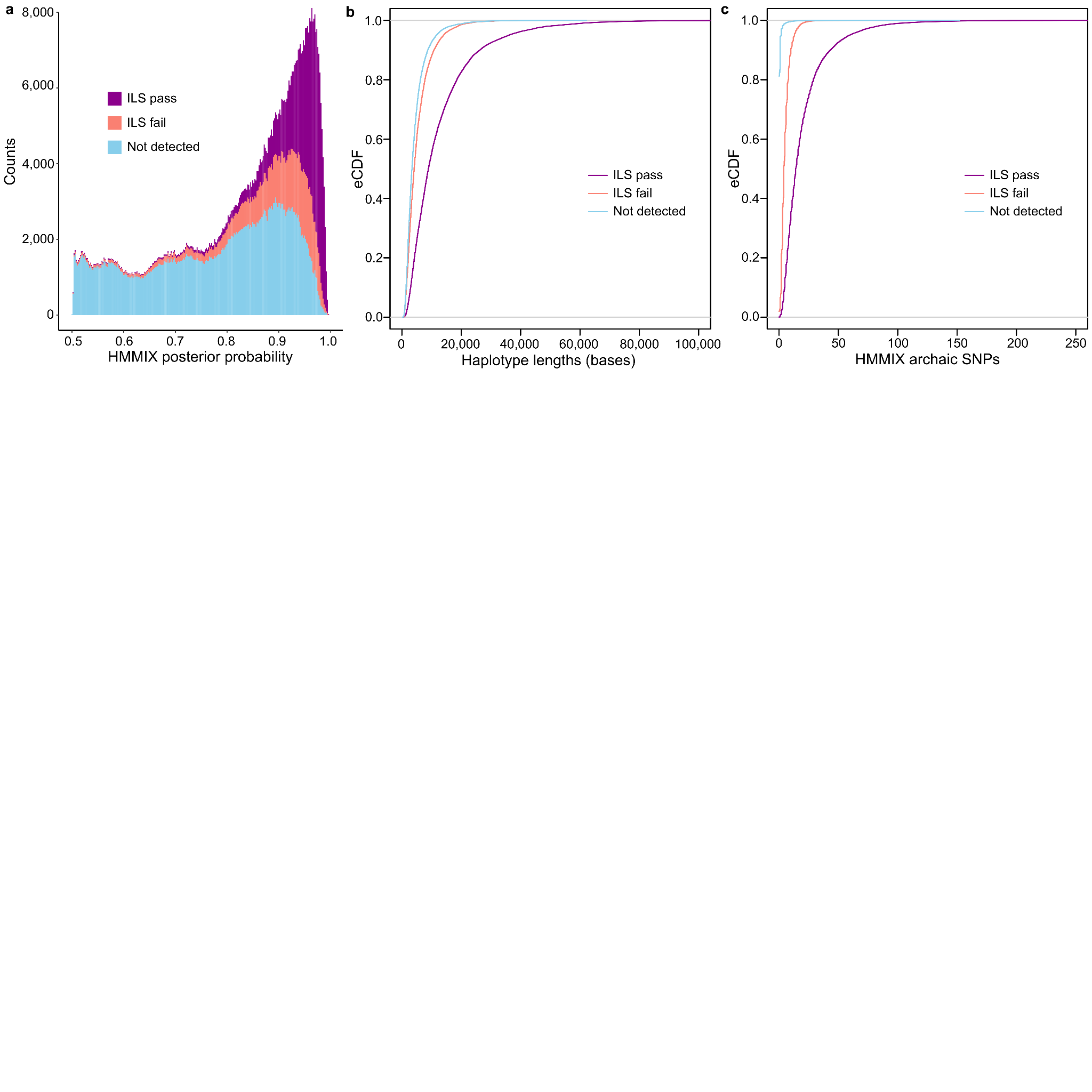

**Figure S1: Evaluation of introgressed segments using HMMIX.**

**(a)** The frequency distribution, depicted as a stacked histogram, represents archaic haplotypes identified in our study, combined with those identified via HMMIX. This panel illustrates the distributions of three distinct haplotype classes: those that either passed the incomplete lineage sorting assessment (ILS pass, purple) or failed to pass (ILS fail, orange), as well as haplotypes solely identified by HMMIX (Not detected, blue). Only haplotypes with a posterior probability greater than 0.5 are shown. **(b-c)** Empirical cumulative density distributions are presented for the lengths (b) and the count of archaic SNPs categorized by HMMIX (c) among the three classes of haplotypes as depicted in panel (a).

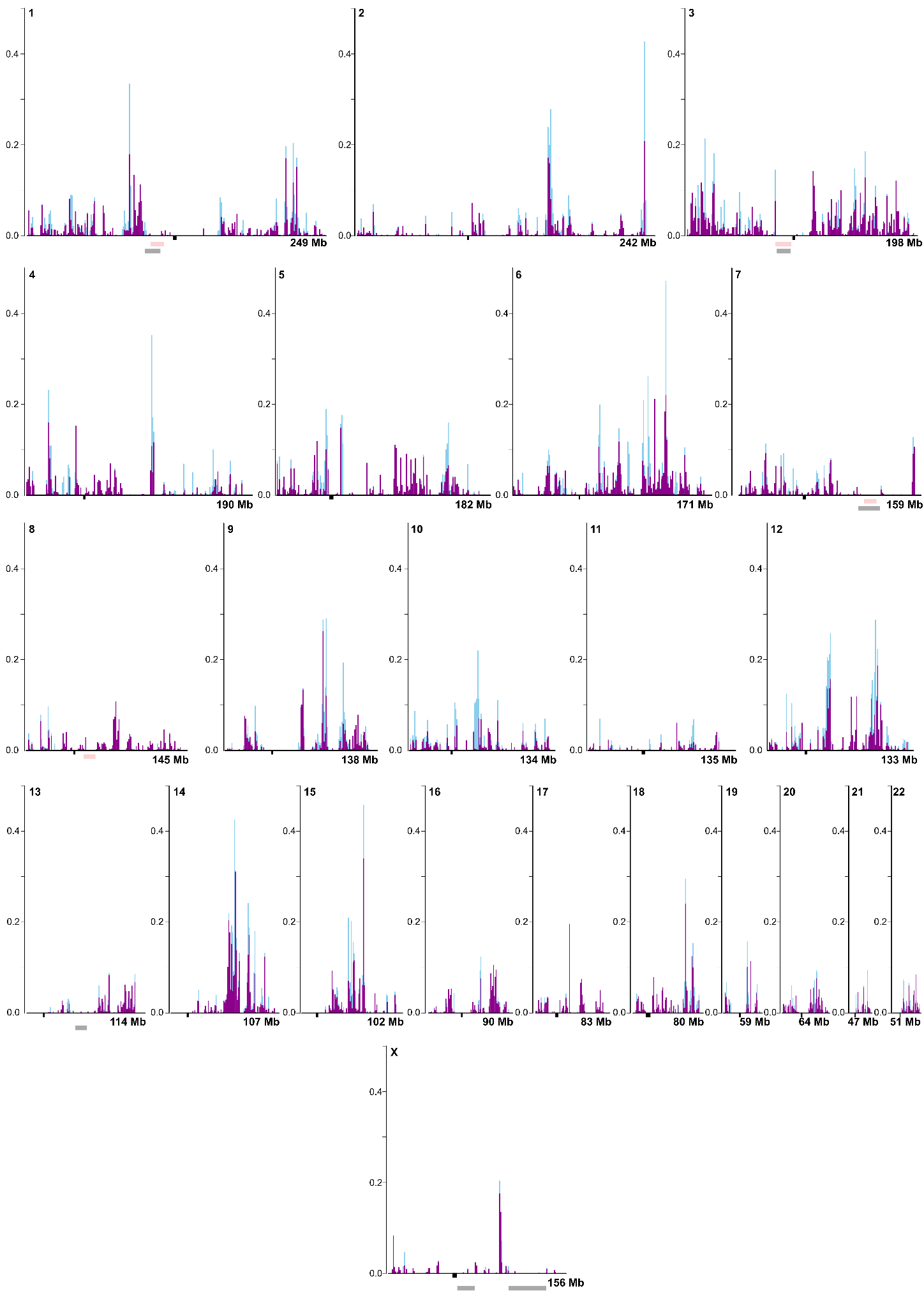

**Figure S2: Genomic distribution of archaic DNA in PNG populations**

The illustration depicts the distribution of archaic DNA fractions, with Neandertal-like sequences represented in blue and Denisovan-like sequences in purple, across 1-megabase windows spanning autosomes and chromosome X. Centromere regions are visualized beneath the x-axis in black. Previously identified genomic regions devoid of archaic DNA are highlighted in gray^31^ and pink^6^.
